# Intraoperative Spinal Cord Stimulation Mitigates Pain after Spine Surgery in Mice

**DOI:** 10.1101/2022.06.09.495484

**Authors:** Satoshi Yamamoto, Alexander Duong, Alex Kim, Chengrui Hu, Blaine Wiemers, Jigong Wang, Jin Mo Chung, Jun-Ho La

**Affiliations:** Department of Anesthesiology, University of Texas Medical Branch, Galveston, TX, USA; Department of Neuroscience and Cell Biology, University of Texas Medical Branch, Galveston, TX, USA

## Abstract

**Background:** Managing postoperative pain after spine surgery is challenging, and up to 40% of operated patients develop failed back surgery syndrome (FBSS) resulting in intractable back and/or leg pain. While spinal cord stimulation (SCS) has been shown to effectively alleviate such chronic pain, it is unknown if intraoperative SCS can mitigate the development of central sensitization that potentially causes intense postoperative pain and FBSS after spine surgery.

**Methods:** As an experimental spine surgery, unilateral T13 laminectomy was performed in mice to expose the dorsal part of L4-5 spinal segments that receive sensory inputs from the hind limb. After the laminectomy, a group of mice received intraoperative SCS epidurally applied to the exposed side of the dorsal part of the spinal cord for an hour under anesthesia before closing the surgical wounds. Secondary mechanical hypersensitivity, a behavioral manifestation of central sensitization, was measured in hind paws using von Frey assay one day before and at predetermined times after surgery. In addition, because von Frey assay is a nocifensive reflex-based analysis that primarily assesses the sensory-discriminative domain of pain, we also performed a conflict avoidance test to capture the affective-motivational domain of pain at selected timepoints post-laminectomy.

**Results:** Mice that underwent unilateral T13 laminectomy developed mechanical hypersensitivity in both hind paws, which gradually resolved in 1-2 weeks. The extent of the hypersensitivity was significantly less in the contralateral hind paw (relative to the laminectomy) than in the ipsilateral hind paw only in females.

Intraoperative SCS applied to the exposed side of the dorsal -spinal cord significantly inhibited the development of hind paw mechanical hypersensitivity only in the SCS-applied side. When paws were mechanically stimulated in their preferred place to present a conflict between pain/discomfort and natural preference, mice avoided the conflict after laminectomy, spending less time in the place than before the surgery. However, mice treated with intraoperative SCS after laminectomy did not avoid the conflict.

**Conclusion:** These results demonstrate that spine surgery for unilateral laminectomy induces central sensitization that results in postoperative pain hypersensitivity.

Intraoperative SCS after laminectomy can mitigate the development of this hypersensitivity in the SCS-applied side.

## Introduction

Low back pain is a very common condition. It is estimated that up to 84% of all adults have low back pain at some point in their lives.^1^ Proportionately, rates of spinal surgical procedures have been rising in the United States.^2^ Nevertheless, 10 to 40 percent of patients develop failed back surgery syndrome (FBSS) following spine surgery.^3–5^ FBSS is a term used to describe back and/or leg pain that persists/recurs or newly develops following one or more operative interventions on the lumbar neuroaxis.^6^ Considering the increasing rates of spine surgery, it is imperative to develop strategies to reduce the risk of developing complications such as FBSS after the surgery.

Surgery can induce central sensitization causing postoperative “pain hypersensitivity,”^7^ and persistent central sensitization is an important underlying contributor to chronic pain states^8^. It is likely that mechanisms of intense postoperative pain after spine surgery involve central sensitization which, if not resolved, may result in FBSS. Therefore, it is an important step toward reducing the risk of FBSS development to prevent spine surgery-induced central sensitization, which translates into preemptive analgesic approaches to reduce postoperative pain after spine surgery. This idea aligns well with the clinical findings that intense postoperative pain itself is a risk factor for pain chronification after surgery and thus adequate postoperative pain management is critical to prevent pain chronification.^9^

Opioid analgesics remain as the mainstay of treatment for postoperative pain. However, many patients have only seen minimal improvement for their postoperative pain management despite these multifaceted efforts.^10,11^

In addition to pharmacological tools, neuromodulation in the form of spinal cord stimulation (SCS) has become an effective option for patients with intractable low back and/or neuropathic leg pain for patients with FBSS.^12,13^ While there is accumulated evidence that SCS is effective for managing chronic pain in FBSS,^14^ it is unknown if SCS can alleviate acute postoperative pain after spine surgery (during which SCS may be temporarily used without permanent device implantation). Considering that intense acute postoperative pain itself is a risk factor for pain chronification after surgery, reduction of postoperative pain after spine surgery may help reduce the risk of FBSS development for reducing or replacing opioid as a countermeasure against the opioid epidemic.^15,16^ Building upon this background, in this study, we first modeled spine surgery-induced pain hypersensitivity in mice and subsequently examined if intraoperative SCS inhibits the development of such hypersensitivity.

## Materials and Methods

### Animals

Adult C57BL/6N mice (9–12 weeks, both sexes, Charles River, Houston, TX) were used throughout this study. Mice were housed with free access to food and water in a temperature, humidity, and light cycle-controlled animal facility accredited by AAALAC (Association for the Assessment and Accreditation of Laboratory and Care International). All procedures and protocols were preregistered and approved by the Institutional Animal Care and Use Committee at the University of Texas Medical Branch and in accordance with the National Institutes of Health (NIH) guidelines. Additional detailed experimental data are available on the Open Science Framework (OSF) (https://osf.io/wpdb9/).^17^ This manuscript adheres to the applicable ARRIVE guidelines.

Mice were randomly stratified into three experimental groups:

1) sham surgery group: 6 females and 7 males
2) laminectomy without SCS group: 8 females and 8 males for the von Frey assay; 5 females and 5 males for the conflict avoidance test
3) laminectomy plus SCS group: 7 females and 7 males for the von Frey assay; 5 females and 5 males for the conflict avoidance test.

### Surgery

Anesthesia was maintained by isoflurane at a concentration of 2%. After the lower back area was shaved, the surgical site was sterilized with povidone and alcohol. A longitudinal midline skin incision was made between the T12 and L1 spinous processes. The lumbosacral fascia was opened longitudinally, and the paravertebral musculature was carefully incised unilaterally in a subperiosteal fashion to expose the left laminae of T12–L1. To expose L4 and L5 spinal segments, the dorsal partial (2 × 1 mm^2^) T13 laminectomy (i.e., laminotomy) on the left lamina was accomplished (Figure 1), and epidural fat was taken out, leaving the dura mater intact and clean. During the surgical procedures, hemostasis was achieved with cotton pads; no cauterization was performed to the surgical site.

**Figure 1.**
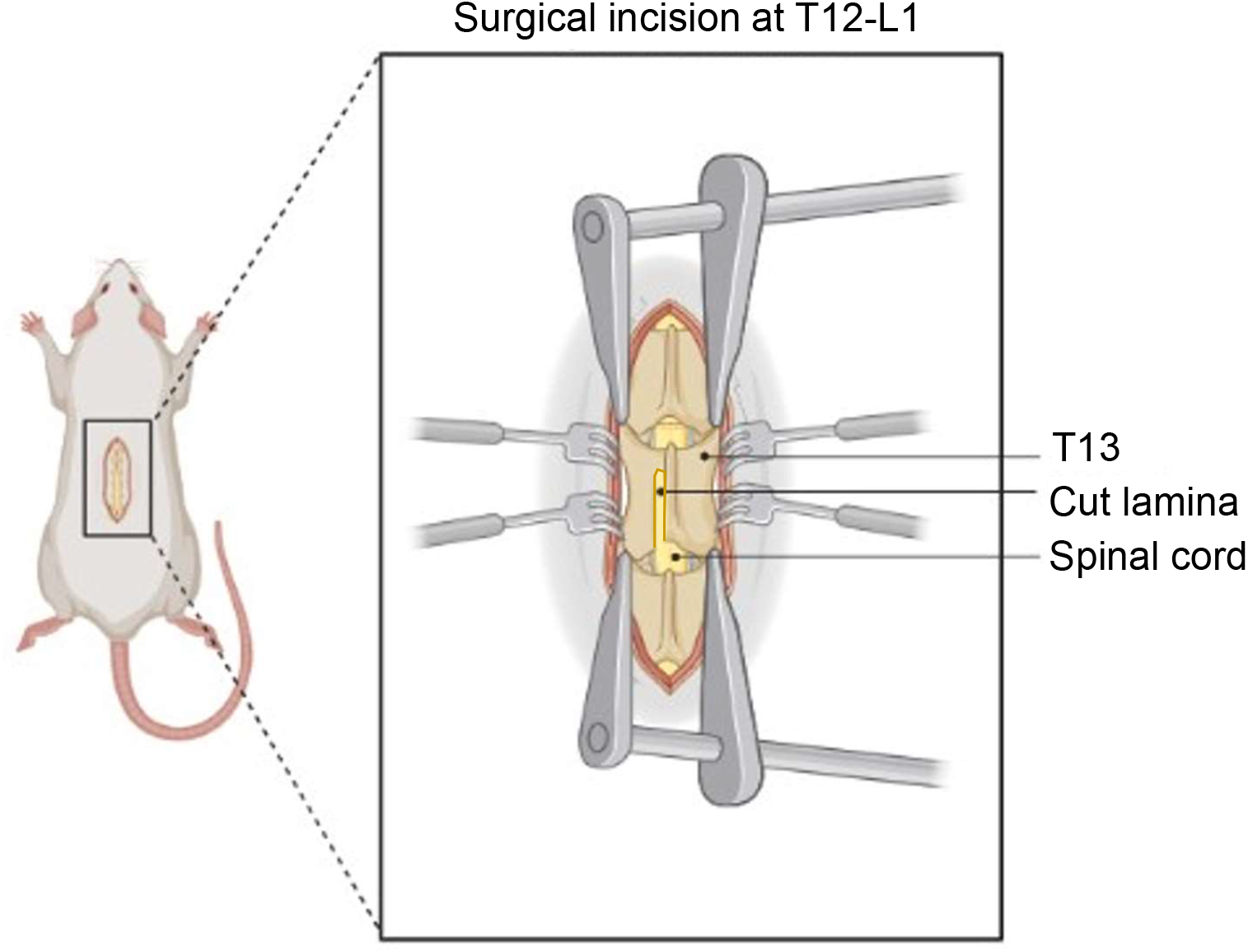
Spine Surgery in a mouse. Partial unilateral laminectomy (i.e., laminotomy) on the left lamina of T13 vertebra was performed. Created with BioRender.com.

After SCS treatment (see below for details) was completed (note that SCS was applied only to the laminectomy plus SCS group), the wounds were closed via propylene suture (Prolene polypropylene sutures; Ethicon; Ethicon Endo-Surgery, Inc., Cincinnati, OH). There were no complications or adverse effects due to materials used. Animals with any neurological deficit in postoperative period were excluded from investigation. All animals were euthanized on postoperative day 14. The sham surgery group underwent the identical surgical procedures excluding the partial laminectomy and SCS. For the laminectomy without SCS group, electrodes were placed epidurally onto the spinal cord without turning on a stimulator.

### Spinal cord stimulation (SCS)

A pair of platinum wire electrode with a ball-shaped round tip were placed epidurally on top of the exposed side of dorsal column and connected to a stimulation isolator (A385 High Current Stimulus Isolator, World Precision Instruments, Sarasota, FL). The isolator received trigger inputs from a pulse stimulator (Model 2100, A-M systems, Sequim, WA) set to deliver biphasic pulses with a 0.2 ms pulse-width at 5 Hz (for detecting motor threshold, MT) or 50 Hz (for SCS). First, MT was determined by gradually increasing the stimulation intensity until contractions of the lower trunk and/or hind limb were visually confirmed. Then, SCS was applied with an amplitude at 50% MT intensity for 60 minutes. We chose these SCS parameters based on the literature showing efficacy in other models.^18–24^ The sham surgery group and laminectomy without SCS group were also kept under anesthesia for 60 minutes before closing surgical wounds.

### Behavior study

Behavioral study was conducted in a dedicated room by experimenters blinded to the nature of experimental groups except for the sex of animals.

#### von Frey assay

Mechanical hypersensitivity was assessed using von Frey filaments. Prior to testing on the day of experiment, mice were placed in the behavioral setup and given 15 minutes to adjust to the setup. Paw withdrawal threshold (PWT) from von Frey filaments was assessed using the “simplified up-down (SUDO) method.”^25^ The PWTs were logarithmically (natural log, ln) transformed to consider Weber’s Law.^26^ Baseline PWTs were determined the day before the surgery (i.e., day -1). The development of postoperative mechanical hypersensitivity was first assessed 120 minutes after the 1-h SCS was turned off on the day of surgery (day 0) and then postoperative days 1, 2, 3, 5, 7, 10 and 14.

#### Conflict avoidance test

Conflict avoidance tests were conducted using a three-compartment apparatus (LE 893, automated place preference system, Panlab/Harvard Apparatus, Barcelona, Spain): Two large chambers (one in white color and the other in black color) were connected by a central grey corridor. Time spent in each chamber was automatically detected by a transducer installed beneath the floor. During a 3-day habituation, mice were allowed to explore all compartments freely for 30□min with smooth floor pads in the two large chambers. Baseline place preference of each mouse was determined on the 3rd day, and mice spending ≥ 2 min in one chamber than in the other were included in this study. At select time points (on day 1 and between days 10-15) after laminectomy, mice were allowed to freely explore the chambers; a bristle floor pad in the preferred chamber and the soft floor pad in the other, causing an emotional-motivational conflict between their natural place preference and pain/discomfort evoked by the mechanical stimulation (i.e., bristles) on their paws. The time spent in the conflict-presented chamber was normalized to that at baseline.

### Data analysis

The simple randomization was conducted between 2 groups (i.e., Sham surgery vs. Laminectomy groups; No SCS vs. SCS groups). The natural log-transformed

PWTs are presented as mean and its 95% confidence. The number of mice for each group is indicated above in the *Animals* section.

We used GraphPad Prism (version 8, GraphPad software, LLC, San Diego, CA, USA) or IBM SPSS Statistics (version 25, SPSS Inc., Chicago, IL, USA) for the 2-way repeated measures ANOVA or mixed-effects analysis followed by Sidak multiple comparison tests between groups at each time point. Specifically, sham surgery group and laminectomy (LMNx) group were compared for each hind paw in each sex (Fig. 2); the degree of LMNx-induced mechanical hypersensitivity was compared either between sexes in each paw or between two paws in each sex (Fig. 3); ‘no SCS’ group and SCS group were compared for each hind paw in each sex (Fig. 4); the degree of SCS-produced inhibition was compared either between sexes in each paw or between two paws in each sex (Fig. 5); ‘no SCS’ group and SCS group were compared in each sex at a given time points (Fig. 6). A p-value (adjusted in case of multiple comparisons) ≤ 0.05 was considered statistically significant. The sample size was estimated using the parameters from preliminary studies for a statistical power of 0.9 at an alpha level 0.01. For example, when male sham and LMNx groups in the contralateral side were compared over multiple time points, 2-way repeated measures ANOVA for the interaction between groups and times (9 time points including the baseline) yielded the Greenhouse-Geisser epsilon of 0.524 and the partial eta squared of 0.2, which estimates at least n=6 per group for the power of 0.9 at the probability of alpha error = 0.01.

**Figure 2.**
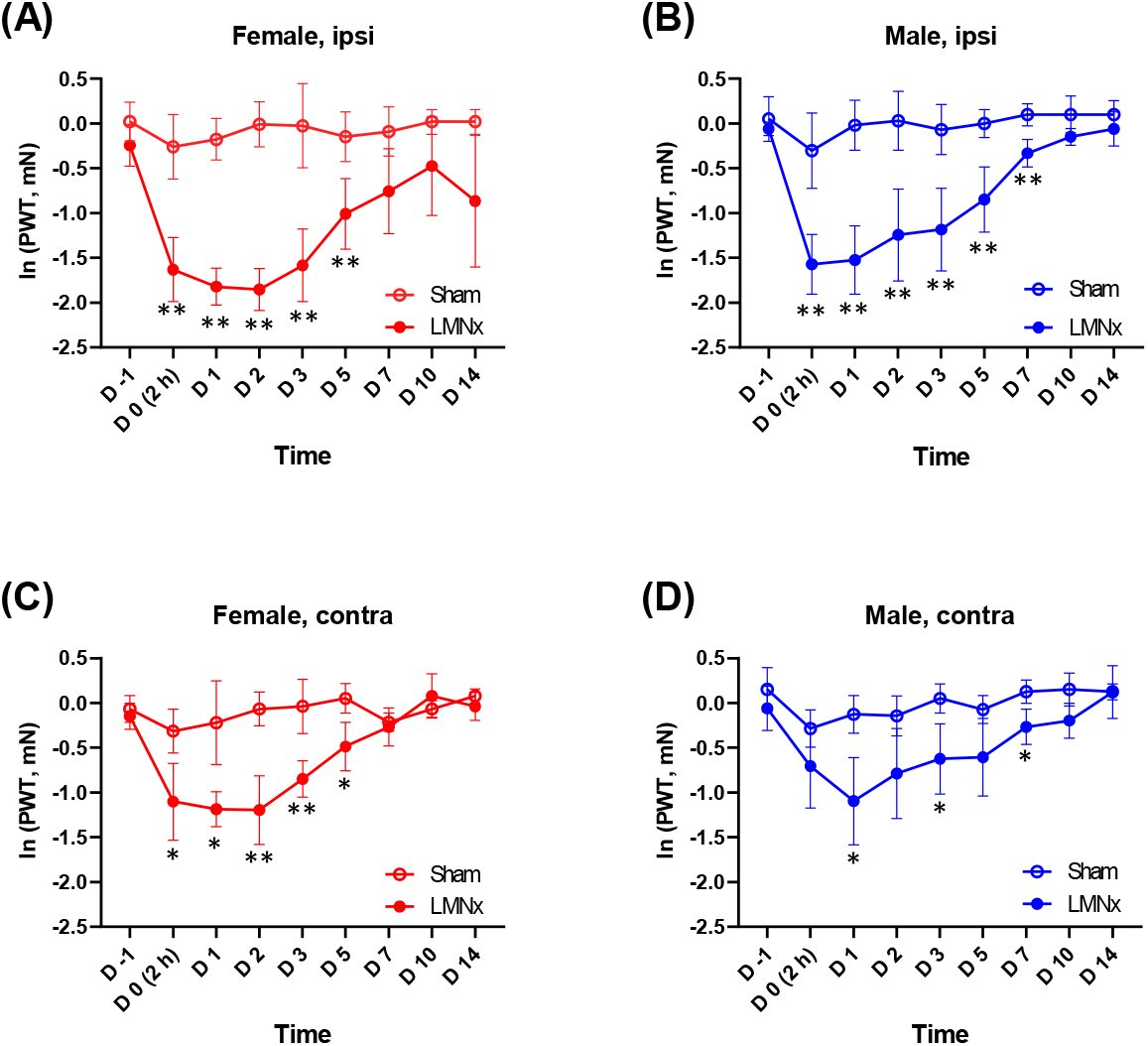
Laminectomy-induced pain hypersensitivity in the ipsilateral and contralateral hind paws. Compared to sham surgery, unilateral laminectomy (LMNx) produced mechanical pain hypersensitivity in the hind paw ipsilateral to the surgery in both (A) females and (B) males. The LMNx also produced mechanical pain hypersensitivity in the contralateral hind paw in (D) females and (E) males. ln (PWT), a natural log-transformed paw withdrawal threshold. *p<0.05 and **p<0.01 by Sidak multiple comparison tests following the mixed effects analysis between Sham and LMNx.

**Figure 3.**
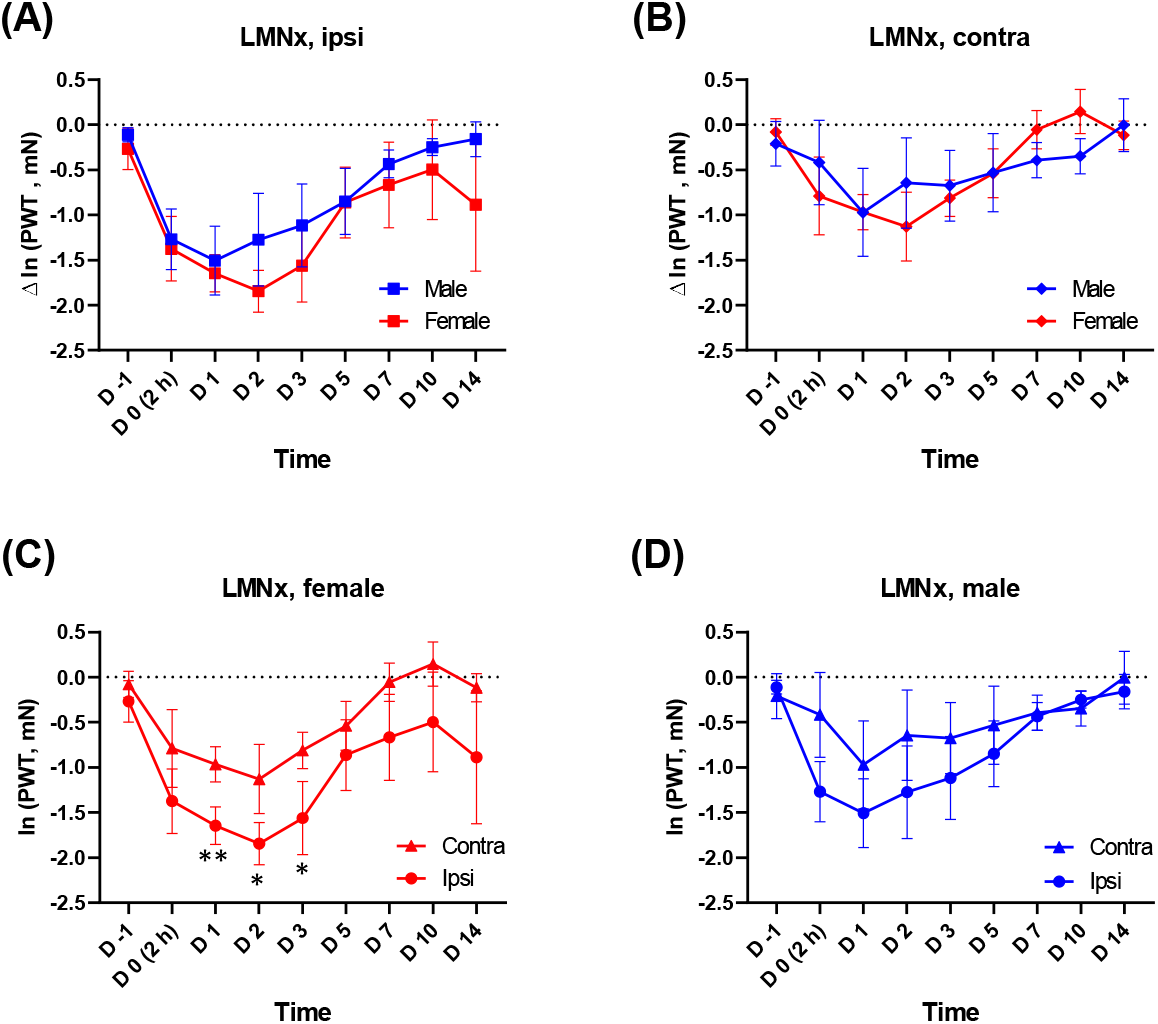
Differences in the degree of laminectomy-induced pain hypersensitivity either between two sexes or between two hind paws. In both (A) ipsi-and (B) contra-lateral hind paws, no sex difference was detected in the degree of LMNx-produced mechanical sensitivity. The degree of the hypersensitivity was significantly greater in the ipsi-than in the contra-lateral side in (C) females, but not in (D) males. *p<0.05 and **p<0.01 by Sidak multiple comparison tests following the mixed effects analysis between Contra and Ipsi. *Δ* ln (PWT), a difference in a natural log-transformed PWT, was calculated by subtracting the mean ln(PWT) of Sham group from individual ln(PWT) in LMNx group at each time points.

**Figure 4.**
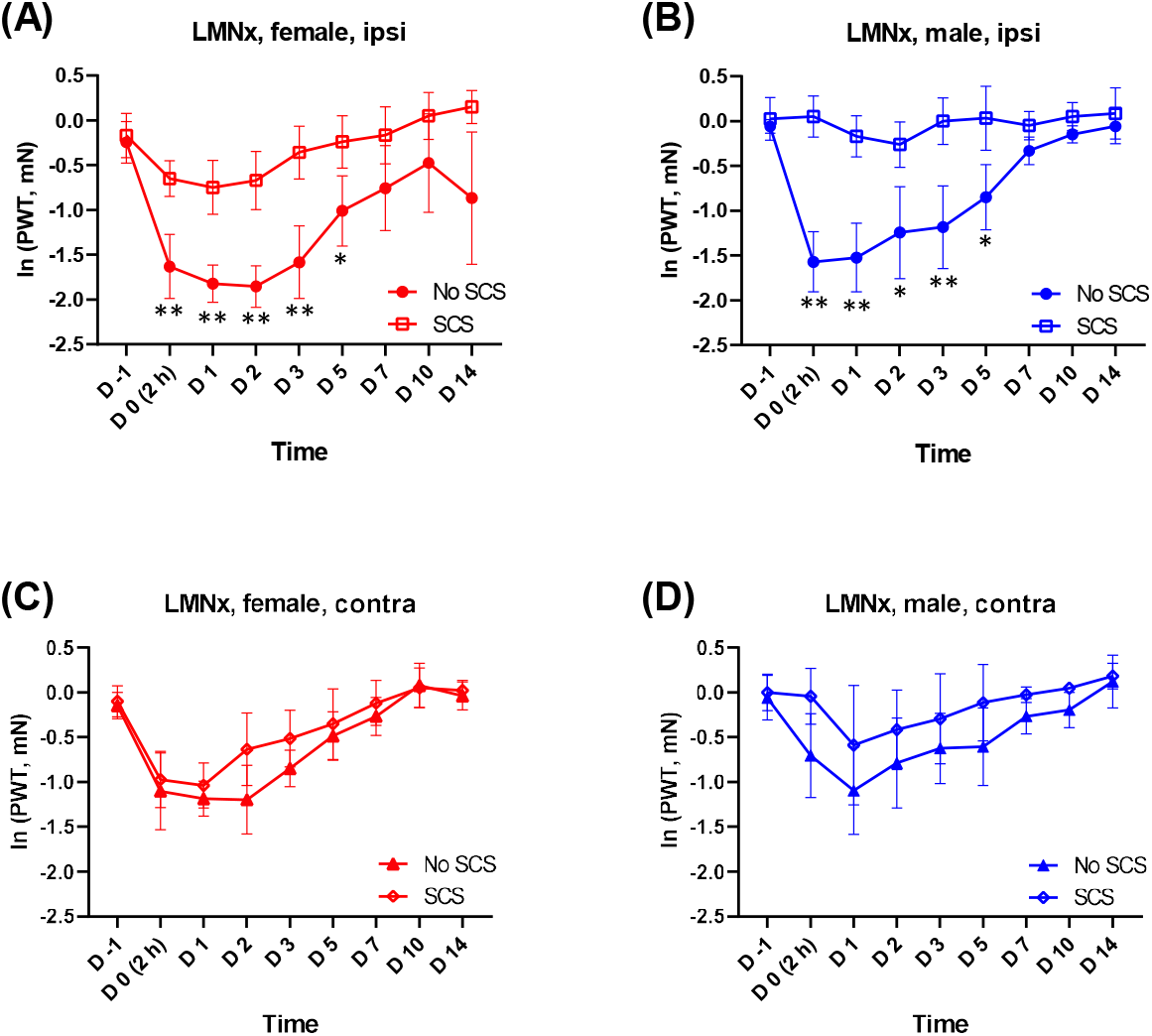
Effect of intraoperative spinal cord stimulation on the laminectomy-induced pain hypersensitivity in the ipsilateral and the contralateral hind paw. In both (A) females and (B) males, unilateral spinal cord stimulation (SCS) after laminectomy (LMNx) mitigated the postoperative mechanical hypersensitivity in the ipsilateral hind paw, but not in the contralateral hind paw (C & D). *p<0.05 and **p<0.01 by Sidak multiple comparison tests following the mixed effects analysis between ‘No SCS’ and SCS groups. Note that ‘No SCS’ group is identical to the LMNx group in Fig. 2.

**Figure 5.**
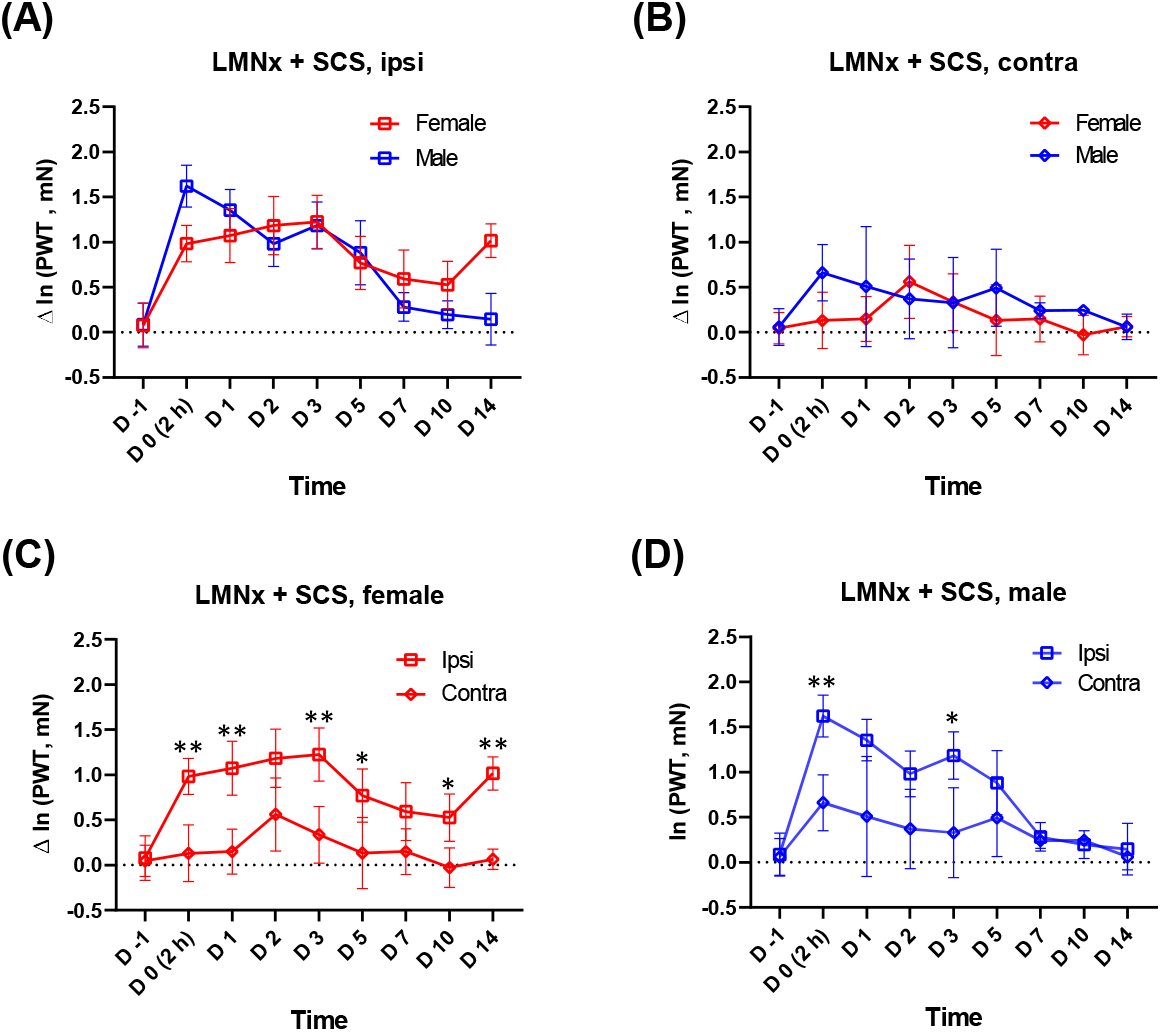
Differences in the degree of intraoperative SCS effects either between two sexes or between two hind paws. In both (A) ipsi-and (B) contra-lateral hind paw, the inhibitory effect of SCS was not different between two sexes. In both (C) females and (D) males, the inhibitory effect of SCS was greater in the ipsi-than in the contra-lateral side. *p<0.05 and **p<0.01 by Sidak multiple comparison test between Ipsi and Contra following the mixed effect analysis. *Δ* ln (PWT), a difference in a natural log-transformed PWT, was calculated by subtracting the mean ln(PWT) of ‘no SCS’ group from individual ln(PWT) in SCS group at each time points.

**Figure 6.**
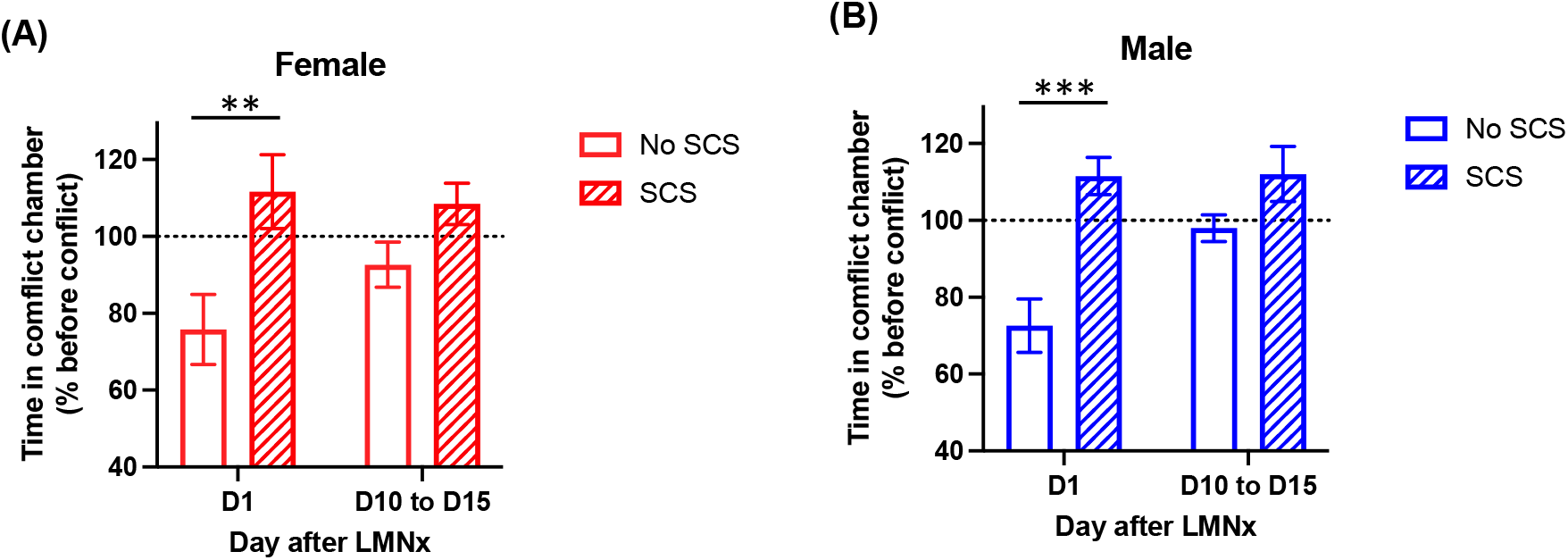
Intraoperative SCS-treated mice do not avoid pain/discomfort-caused conflict after laminectomy. Conflict avoidance test results between unilateral laminectomy (LMNx)-treated (A) female and (B) male mice with and without intraoperative spinal cord stimulation (SCS). X-axis represents the days post-operation. Y-Axis represents the % of time that a mouse spent in the preferred chamber despite pain/discomfort-caused conflict. **p<0.01, ***p<0.001 vs. corresponding ‘No SCS’ group.

## Results

### Unilateral T13 laminectomy produces bilateral mechanical hypersensitivity in hind paws

We performed a unilateral T13 laminectomy to expose the dorsal part of L4-5 spinal segments^27^ to which peripheral sensory fibers innervating a hind limb centrally project. As shown in Figure 2, compared to sham surgery group, laminectomy (LMNx) group developed postoperative mechanical hypersensitivity in the hind paw ipsilateral to the surgery side, showing significantly reduced withdrawal thresholds (Fig. 2A: F(1,12)=36.1, p<0.01 in females; Fig. 2B: F(1,13)=41.3, p<0.01 in males). In both sexes, there was a significant interaction between groups (sham vs. LMNx) and times (F(8,96)=12.4, p<0.01 in females; F(8,104)=15.7, p<0.01 in males), showing that this hypersensitivity gradually resolved in 7-14 days after LMNx. In the contralateral hind paw, LMNx also resulted in mechanical hypersensitivity (Fig. 2C: F(1,12)=35.6, p<0.01 in females; Fig. 2D: F(1,13)=13.4, p<0.01 in males) which subsided over times similarly to that in the ipsilateral hind paw.

To delineate the degree of “LMNx-induced” pain hypersensitivity and compare the degree either between sexes in each hind paw or between hind paws in each sex, we subtracted the mean PWT of sham surgery group from PWTs of individual mice in LMNx group at each time point (difference PWT, *Δ*) and ran a statistical test for the four datasets (2 sexes x 2 hind paws: F(3,241)=5.41, p<0.01). Subsequent multiple comparison tests revealed that there was no sex difference in the degree of LMNx-induced mechanical hypersensitivity in either ipsi-or contra-lateral hind paws (Fig. 3A and B). However, in females only, LMNx induced mechanical hypersensitivity to a lesser degree in the contralateral hind paw (Fig. 3C: t(241)=3.54, Sidak-adjusted p<0.01).

### Intraoperative unilateral SCS mitigates the development of hind paw mechanical hypersensitivity after laminectomy

Next, we determined if intraoperative SCS, applied to the exposed side of L4-5 dorsal spinal cord, would affect the development of LMNx-induced mechanical hypersensitivity in hind paws. As shown in Figure 4, intraoperative SCS significantly reduced the LMNx-induced mechanical hypersensitivity in the ipsilateral hind paw in both sexes (Fig. 4A: F(1,13)=23.8, p<0.01 in females; Fig. 4B: F(1,13)=39.1, p<0.01 in males). By contrast, in the contralateral hind paw, there was no significant inhibitory effect of SCS on LMNx-induced mechanical hypersensitivity (Fig. 4C & D), indicating that intraoperative SCS on the exposed side of dorsal column mitigate LMNx-induced mechanical hypersensitivity only in the SCS-applied side.

We compared the degree of “SCS-produced” inhibition of postoperative mechanical hypersensitivity either between sexes in each hind paw or between hind paws in each sex. To this end, we subtracted the mean PWT of ‘No SCS’ group from PWTs of individual mice in SCS group at each time point (difference PWT, *Δ*) and ran a statistical test for the four datasets (2 sexes x 2 hind paws:

F(3,199)=15.3, p<0.01). Subsequent multiple comparison tests revealed that there was no sex difference in the efficacy of SCS in both hind paws (Fig. 5A and B). However, as expected by the previous results in Fig. 4C and D, an increase in PWT by SCS was significantly greater in the ipsi-than in the contra-lateral hind paw in both sexes (Fig. 5C: t(199)=5.59, p<0.01 in females; Fig. 5D: t(199)=3.81, p<0.01 in males).

### Intraoperative SCS-treated mice did not avoid pain/discomfort-caused conflict after laminectomy

We further assessed affective-motivational aspect of pain due to postoperative mechanical hypersensitivity. One day after LMNx, both female and male ‘No SCS’ groups avoided the conflict-presented chamber (i.e., the chamber of natural preference but with mechanical stimulation by floor bristles), spending less time in the chamber than at baseline. By contrast, SCS-treated group did not avoid the conflict, spending significantly more time than their counterparts (Fig. 5A: t(16)=3.284, p<0.01 in females; Fig. 5B: t(14)=4.914, p<0.001 in males). Between days 10-15 after LMNx, when LMNx-induced mechanical hypersensitivity was found to be resolved in the von Frey assay (Fig. 2), ‘No SCS’ group no longer avoided the conflict-presented chamber.

## Discussion

Despite unilateral partial laminectomy (i.e., laminotomy) being the least invasive type of procedure among spine surgeries, the present study demonstrates that mice develop bilateral postoperative mechanical hypersensitivity in hind paws after this procedure, suggesting that laminectomy induces central sensitization that produces pain hypersensitivity even in the body area (both hind paws) remote from the surgical injury site (the back) and in the injury-associated dermatome (the ipsilateral hind paw). By contrast, sham surgery did not produce any obvious mechanical hypersensitivity in hind paws, indicating that soft tissue injury alone in the back, even though deep enough to expose the vertebrae, is not sufficiently intense to induce central sensitization resulting in hind paw mechanical hypersensitivity. However, it should be mentioned that after the soft tissue injury by sham surgery, postoperative pain hypersensitivity may still develop at/around the surgical wounds in the back.

The results of conflict avoidance tests in this study indicate that mice experience pain/discomfort upon mechanical stimulation of (hind) paws after laminectomy to the extent that their natural place preference is diminished. Importantly, mice treated with intraoperative SCS after laminectomy did not show a decrease in their place preference despite the mechanical stimulation, suggesting that these mice experienced less pain/discomfort than their counterparts because the SCS had mitigated the development of central sensitization and resultant mechanical hypersensitivity. This notion is supported by the observation that when mechanical hypersensitivity has resolved (between days 10-15), operated mice without SCS treatment no longer showed conflict avoidance.

In clinics, surgical interventions are considered after multiple trials of conservative, nonsurgical approaches for patients suffering back pain due to anatomical spinal nerve compression. Decompressive laminectomy, with or without fusion, is the most common surgery for spinal stenosis and symptomatic degenerative spondylolisthesis, which involves removal of a part/all of one lamina or both laminae of the vertebra at the affected level. After spine surgeries, postoperative pain control is a challenge.^28^ In this regard, the present findings that a lumbar spinal cord-exposing laminectomy caused bilateral mechanical hypersensitivity in hind paws may be a preclinical mirror of intense postoperative pain and/or the new onset of leg pain after spine surgery in patients.

In the context of prevention of chronic surgical pain, severe acute postoperative pain itself is known as a risk factor for pain chronification,^29^ and thus, it has become apparent that preemptive analgesia is a useful approach beyond its expected duration. Since the spinal cord becomes accessible for SCS after laminectomy, intraoperative SCS can be utilized as a preemptive analgesic approach for reducing postoperative pain after spine surgery. Our present preclinical study clearly demonstrates the efficacy of intraoperative SCS against hind paw mechanical hypersensitivity development after laminectomy, suggesting that this preemptive approach is a feasible and promising option to mitigate postoperative pain after spine surgery, which may further reduce the risk of FBSS development. SCS has shown its effectiveness on chronic pain in patients with FBSS.^30–34^ Although there is a report that a neuromodulation technique has a strong positive effect on acute postoperative pain,^35^ to the best of our knowledge, the intraoperative use of SCS is the novel strategy to control acute postoperative pain after spine surgery.

The mechanisms of SCS are still unclear. Some propose both orthodromic and antidromic activation of large myelinated fibers in the dorsal column, recruitment of supraspinal nuclei to bolster descending inhibitory impulses, and changed cerebrospinal fluid (CSF) neurochemistry through controlled delivery of electrical impulses to the dorsal columns in the spinal cord.^36^ It should be mentioned that in this study, one-time intraoperative session has a long-lasting effect on the development of postoperative pain hypersensitivity. This suggests that intraoperative SCS counteracts the mechanism(s) of long-term central sensitization, so that the sensitization does not fully develop. Future research is warranted to elucidate how SCS reins on the central sensitization development and if different SCS parameters would produce better outcomes (e.g., also effective on central sensitization in the contralateral side) than what we observed in this study using SCS with 0.2 ms pulses at 50 Hz frequency and 50% MT intensity.

Altogether, our findings suggest that spine surgery poses a significant risk for intense postoperative pain due to central sensitization. Intraoperative SCS can mitigate the development of laminectomy-induced central sensitization, being a feasible and promising option as a preemptive analgesic approach to reduce postoperative pain after spine surgery and reduce the risk of FBSS development. Further investigation is needed to understand the mechanisms of SCS and SCS parameters for optimal outcomes.

## References

1. Chan C, Peng P. Failed Back Surgery Syndrome. Pain Med 2011;12:577–606.

2. Gray DT, Deyo RA, Kreuter W, Mirza SK, Heagerty PJ, Comstock BA, Chan L. Population-based trends in volumes and rates of ambulatory lumbar spine surgery. Spine 2006;31:1957–63; discussion 1964.

3. Wilkinson HA. The Failed Back Syndrome: Etiology and Therapy. Springer Science & Business Media, 2012.

4. Lehmann TR, LaRocca HS. Repeat lumbar surgery. A review of patients with failure from previous lumbar surgery treated by spinal canal exploration and lumbar spinal fusion. Spine 1981;6:615–9.

5. Law JD, Lehman RA, Kirsch WM. Reoperation after lumbar intervertebral disc surgery. J Neurosurg 1978;48:259–63.

6. Thomson S. Failed back surgery syndrome – definition, epidemiology and demographics. Br J Pain 2013;7:56–9.

7. Pak DJ, Yong RJ, Kaye AD, Urman RD. Chronification of Pain: Mechanisms, Current Understanding, and Clinical Implications. Curr Pain Headache Rep 2018;22:9.

8. Hankerd K, McDonough KE, Wang J, Tang S-J, Chung JM, La J-H. Postinjury stimulation triggers a transition to nociplastic pain in mice. Pain 2021.

9. McGreevy K, Bottros MM, Raja SN. Preventing Chronic Pain following Acute Pain: Risk Factors, Preventive Strategies, and their Efficacy. Eur J Pain Suppl 2011;5:365– 72.

10. Gordon DB, Leon-Casasola OA de, Wu CL, Sluka KA, Brennan TJ, Chou R. Research Gaps in Practice Guidelines for Acute Postoperative Pain Management in Adults: Findings From a Review of the Evidence for an American Pain Society Clinical Practice Guideline. J Pain 2016;17:158–66.

11. Gan TJ, Habib AS, Miller TE, White W, Apfelbaum JL. Incidence, patient satisfaction, and perceptions of post-surgical pain: results from a US national survey. Curr Med Res Opin 2014;30:149–60.

12. Sdrulla A, Guan Y, Raja S. Spinal Cord Stimulation: Clinical Efficacy and Potential Mechanisms. Pain Pract Off J World Inst Pain 2018;18:1048–67.

13. Labaran L, Jain N, Puvanesarajah V, Jain A, Buchholz AL, Hassanzadeh H. A Retrospective Database Review of the Indications, Complications, and Incidence of Subsequent Spine Surgery in 12,297 Spinal Cord Stimulator Patients. Neuromodulation J Int Neuromodulation Soc 2019.

14. Taylor RS, Van Buyten J-P, Buchser E. Spinal cord stimulation for chronic back and leg pain and failed back surgery syndrome: a systematic review and analysis of prognostic factors. Spine 2005;30:152–60.

15. Bonnie RJ, Kesselheim AS, Clark DJ. Both Urgency and Balance Needed in Addressing Opioid Epidemic: A Report From the National Academies of Sciences, Engineering, and Medicine. JAMA 2017;318:423–4.

16. Manchikanti L, Helm S, Fellows B, Janata JW, Pampati V, Grider JS, Boswell MV. Opioid epidemic in the United States. Pain Physician 2012;15:ES9–38.

17. Yamamoto S. Intraoperative Spinal Cord Stimulation Mitigates Pain after Spine Surgery in Mice. 2022. Available at: https://osf.io/wpdb9/. Accessed May 8, 2022.

18. Song Z, Meyerson BA, Linderoth B. High-Frequency (1lJkHz) Spinal Cord Stimulation-Is Pulse Shape Crucial for the Efficacy? A Pilot Study. Neuromodulation J Int Neuromodulation Soc 2015;18:714–20.

19. Song Z, Viisanen H, Meyerson BA, Pertovaara A, Linderoth B. Efficacy of kilohertz-frequency and conventional spinal cord stimulation in rat models of different pain conditions. Neuromodulation J Int Neuromodulation Soc 2014;17:226–34; discussion 234-235.

20. Meyerson BA, Ren B, Herregodts P, Linderoth B. Spinal cord stimulation in animal models of mononeuropathy: effects on the withdrawal response and the flexor reflex. Pain 1995;61:229–43.

21. Shechter R, Yang F, Xu Q, Cheong Y-K, He S-Q, Sdrulla A, Carteret AF, Wacnik PW, Dong X, Meyer RA, Raja SN, Guan Y. Conventional and kilohertz-frequency spinal cord stimulation produces intensity-and frequency-dependent inhibition of mechanical hypersensitivity in a rat model of neuropathic pain. Anesthesiology 2013;119:422–32.

22. Sato KL, Johanek LM, Sanada LS, Sluka KA. Spinal cord stimulation reduces mechanical hyperalgesia and glial cell activation in animals with neuropathic pain. Anesth Analg 2014;118:464–72.

23. Beek M van, Kleef M van, Linderoth B, Kuijk SMJ van, Honig WM, Joosten EA. Spinal cord stimulation in experimental chronic painful diabetic polyneuropathy: Delayed effect of High-frequency stimulation. Eur J Pain Lond Engl 2017;21:795–803.

24. Meuwissen KPV, Gu JW, Zhang TC, Joosten EAJ. Conventional-SCS vs. Burst-SCS and the Behavioral Effect on Mechanical Hypersensitivity in a Rat Model of Chronic Neuropathic Pain: Effect of Amplitude. Neuromodulation J Int Neuromodulation Soc 2018;21:19–30.

25. Bonin RP, Bories C, De Koninck Y. A simplified up-down method (SUDO) for measuring mechanical nociception in rodents using von Frey filaments. Mol Pain 2014;10:1744–8069.

26. Mills C, Leblond D, Joshi S, Zhu C, Hsieh G, Jacobson P, Meyer M, Decker M. Estimating efficacy and drug ED50’s using von Frey thresholds: impact of weber’s law and log transformation. J Pain 2012;13:519–23.

27. Harrison M, O’Brien A, Adams L, Cowin G, Ruitenberg MJ, Sengul G, Watson C. Vertebral landmarks for the identification of spinal cord segments in the mouse. NeuroImage 2013;68:22–9.

28. Maheshwari K, Avitsian R, Sessler DI, Makarova N, Tanios M, Raza S, Traul D, Rajan S, Manlapaz M, Machado S, Krishnaney A, Machado A, Rosenquist R, Kurz A. Multimodal Analgesic Regimen for Spine Surgery A Randomized Placebo-controlled Trial. Anesthesiology 2020;132:992–1002.

29. Aasvang EK, Brandsborg B, Christensen B, Jensen TS, Kehlet H. Neurophysiological characterization of postherniotomy pain. Pain 2008;137:173–81.

30. Zucco F, Ciampichini R, Lavano A, Costantini A, De Rose M, Poli P, Fortini G, Demartini L, De Simone E, Menardo V, Cisotto P, Meglio M, Scalone L, Mantovani LG. Cost-Effectiveness and Cost-Utility Analysis of Spinal Cord Stimulation in Patients With Failed Back Surgery Syndrome: Results From the PRECISE Study. Neuromodulation J Int Neuromodulation Soc 2015;18:266–76; discussion 276.

31. Rigoard P, Jacques L, Delmotte A, Poon K, Munson R, Monlezun O, Roulaud M, Prevost A, Guetarni F, Bataille B, Kumar K. An algorithmic programming approach for back pain symptoms in failed back surgery syndrome using spinal cord stimulation with a multicolumn surgically implanted epidural lead: a multicenter international prospective study. Pain Pract Off J World Inst Pain 2015;15:195–207.

32. Remacle TY, Bonhomme VL, Renwart H-JP, Remacle JM. Effect of Multicolumn Lead Spinal Cord Stimulation on Low Back Pain in Failed Back Surgery Patients: A Three-Year Follow-Up. Neuromodulation J Int Neuromodulation Soc 2017;20:668–74.

33. Paul AR, Kumar V, Roth S, Gooch MR, Pilitsis JG. Establishing Minimal Clinically Important Difference of Spinal Cord Stimulation Therapy in Post-Laminectomy Syndrome. Neurosurgery 2017;81:1011–5.

34. Sweet J, Badjatiya A, Tan D, Miller J. Paresthesia-free high-density spinal cord stimulation for postlaminectomy syndrome in a prescreened population: a prospective case series. Neuromodulation Technol Neural Interface 2016;19:260–7.

35. Lawson McLean A, Kalff R, Reichart R. Spinal Cord Stimulation for Acute Pain Following Surgery for Cervical Myelopathy: A Novel Treatment Strategy. Pain Pract Off J World Inst Pain 2019;19:310–5.

36. Oakley JC, Prager JP. Spinal cord stimulation: mechanisms of action. Spine 2002;27:2574–83.

